# A human supergene: *BRCA1, CCDC200*, and U2 snRNAs

**DOI:** 10.1101/2025.11.08.687349

**Authors:** Pelle Scholten, Michael Segel, David Haig

## Abstract

*BRCA1* occurs on one of two major haplotypes in populations of European and Asian ancestry that show signs of selection and functional differences. Near-maximal linkage disequilibrium associated with these haplotypes extends for more than 300 kilobases from *RND2* to *RNU2*. There is greater haplotypic diversity in populations from Africa, with the majority haplotype in Europe and Asia a minority haplotype in Africa. Recombination across this region is very rare or absent, both within Africa and outside of Africa, with haplotypes inherited as ‘alleles’ of a supergene. The polymorphism appears to be maintained by some form of balancing selection. We analyzed the repeat structure of the *RND2–RNU2* supergene using the CHM13 telomere-to-telomere genome assembly. The most notable feature is the highly regular array of 6.1-kb *RNU2* repeats. Immediately adjacent to the *RNU2* array are retroviral sequences that may play a role in the concerted evolution of the repeats. Other structure-forming sequences from the *RND2– RNU2* region are predicted to be sources of replicative stress and may interfere with meiotic chromosome synapsis.

## Introduction

Soon after its identification as a gene mutated in familial breast and ovarian cancer (Miki et al. 1994), *BRCA1* was found to reside in an unusual part of the genome. Almost complete linkage disequilibrium extended from *RND2* to *RNU2* in European and Asian populations, encompassing the entire *BRCA1* coding sequence (Liu and Barker 1999). Chromosomes could be assigned to one of two major haplotypes that behaved as if they were alleles at a single Mendelian locus. Meiotic recombination appeared to be suppressed both in haplotypic heterozygotes and homozygotes because rare variants were restricted to one or other haplotype and were in maximal linkage disequilibrium with respect to each other within haplogroups. The *RNU2* locus contained a tandem array of 6.1-kb repeats, each associated with the coding sequence of a U2 snRNA. These repeats maintained greater than 99% identity within arrays while preserving linkage disequilibrium between sequences on opposite flanks of the array, consistent with a recent history of intrachromosomal exchanges but rare interchromosomal exchanges (Liao et al. 1997).

Despite intense subsequent interest in *BRCA1*, its unusual genomic milieu has attracted little attention, although the extreme rarity of meiotic recombination between *RNU2* and *BRCA1* has been confirmed in a study of distantly related families carrying the same *BRCA1* founder mutation (Tessereau et al. 2014). In this paper, we first characterize worldwide haplotypes of *BRCA1–RNU2* and confirm Liu and Barker’s (1999) observation of two major haplotypes in Eurasian populations. Greater haplotypic diversity exists in Africa, but long-range linkage disequilibrium is maintained. We conclude that the *BRCA1–RNU2* region satisfies criteria to be considered a ‘supergene’ with the two haplogroups maintained by some form of balancing selection. We then characterize the genomic structure of the region of suppressed recombination and find that it contains multiple sequences predicted to form strong secondary structures that are likely to be causes of replicative stress. We conjecture that these structures also contribute to the suppression of recombination by interfering with meiotic synapsis. Finally, we find that *RNU2* is associated with retroviral coding sequences that may be involved in the concerted evolution of the *RNU2* tandem array. We conclude by considering what supergenic phenotypes may be under selection.

### Haplotypic diversity

*BRCA1* haplotypes can be tagged by four common SNPs that cause amino-acid substitutions in BRCA1 protein (Table 1). Three of these SNPs (rs799917, rs16941, rs16942) occur within a 937 nucleotide segment of exon 11 (a very large internal exon) with the fourth SNP (rs1799966) located 21 kb distant in exon 16.

**Table 1.** Common non-synonymous SNPs in the *BRCA1* coding sequence.

| SNP | Ancestral allele | Derived allele |
| --- | --- | --- |
| rs799917 | T; Leu831 (L) | C; Pro831 (P) |
| rs16941 | A; Glu1038 (E) | G; Gly1038 (G) |
| rs16942 | G; Arg1183 (R) | A; Lys1183 (K) |
| rs1799966 | A; Ser1613 (S) | G; Gly1613 (G) |

SNPs in Table 1 are identified by the nucleotide on the template strand and by encoded amino acid. The standard one-letter symbols for amino acids (bracketed letters) will be used for short-hand descriptions of haplotypes (e.g.

LEKS, PEKS, LGRG). Ancestral alleles were identified as the variant present in the reference sequences of chimpanzees, bonobos, and gorillas. Twenty-five chimpanzees and thirteen bonobos all possessed LERS haplotypes but LERS was not found in our sample of 2503 modern human individuals.

As first reported by Liu and Barker (1999), populations outside of Africa are dominated by two major haplotypes. Two of the derived alleles (Gly1038, Gly1613) occur on LGRG haplotypes and two (Pro831, Lys1183) occur on PEKS haplotypes. All four SNPs are in near-maximal linkage disequilibrium with each other and with many other SNPs between *RND2* and *RNU2*. The region of near-maximal linkage disequilibrium extends for more than 330 kilobases on the GRCh38 assembly (Schneider et al. 2013) and for more than 370 kilobases on the CHM13 telomere-to-telomere assembly (Nurk et al. 2022). The GRCh38 assembly is a composite of clones. It possesses the majority European alleles (PEKS) for the four-tagging SNPs. CHM13 contains an LGRG haplotype. Intronic sequences of *BRCA1* are more than 99% identical between GRCh38 and CHM13.

In a principal-components analysis of African haplotypes based on a large number of SNPs (Figure 1a), the first principal component (PC1) separated LGRG and LERG haplotypes (negative values on PC1) from PEKS and LEKS haplotypes (positive values on PC1). The second principal component (PC2) captured variation among PEKS and LEKS haplotypes. All PEKS haplotypes, and a subset of LEKS haplotypes, had negative values on PC1 with positive values on PC2. The remainder of the LEKS haplotypes had negative values on PC1 and PC2. In a similar principal-components analysis of non-African haplotypes (Figure 1b), PC1 explained 87% of the variance and separated LGRG haplotypes from PEKS haplotypes with PC2 explaining only 2% of the variance (capturing variation among LGRG haplotypes). Although PEKS is the majority haplotype in Figure 1b, this is not obvious from the Figure: PEKS haplotypes are piled on top of each other because of their high genetic similarity. Figures 1a and 1b show that the four tagging SNPs of Table 1 are successful in distinguishing between major classes of haplotypes.

**Figure 1.**
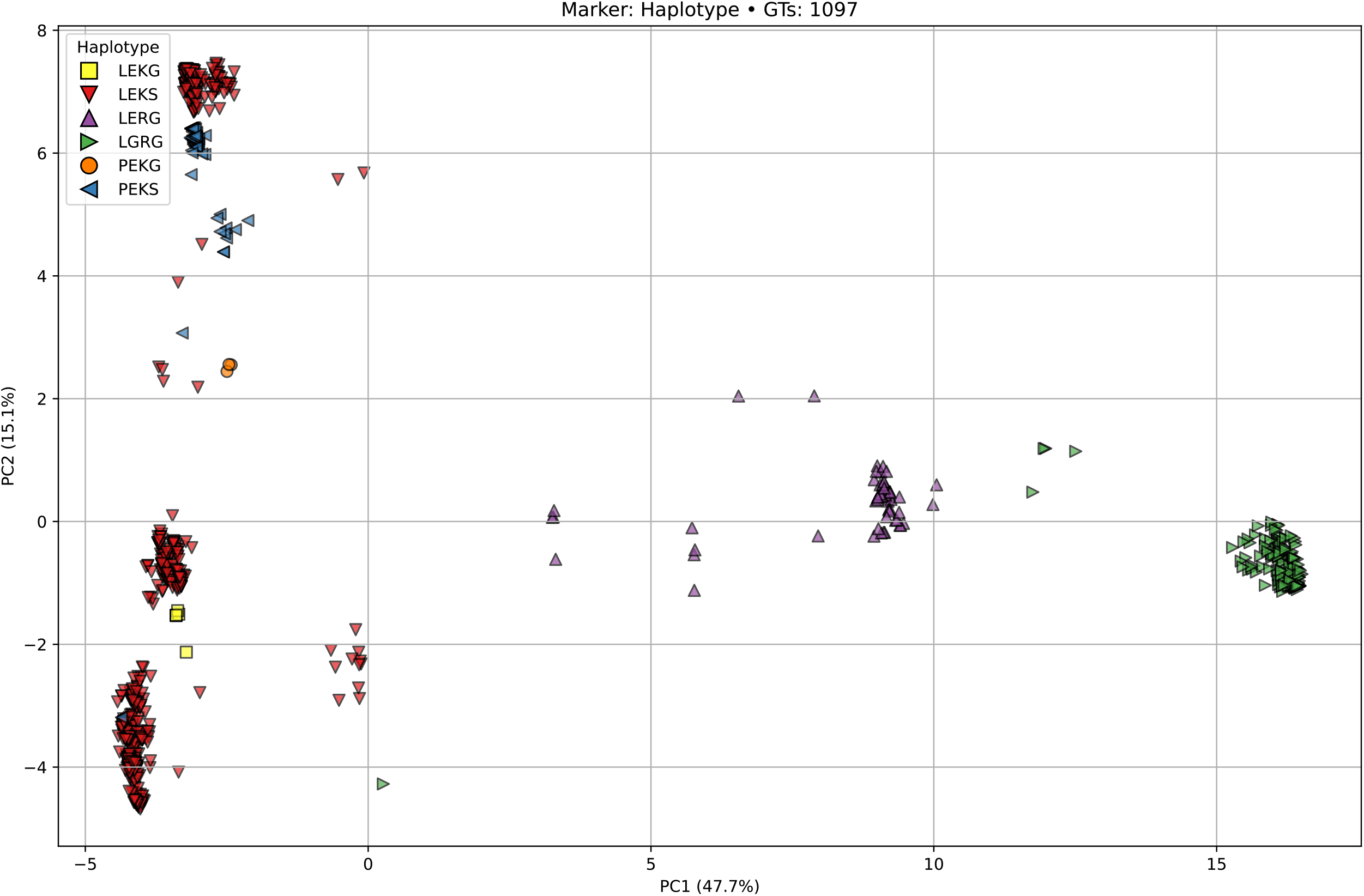
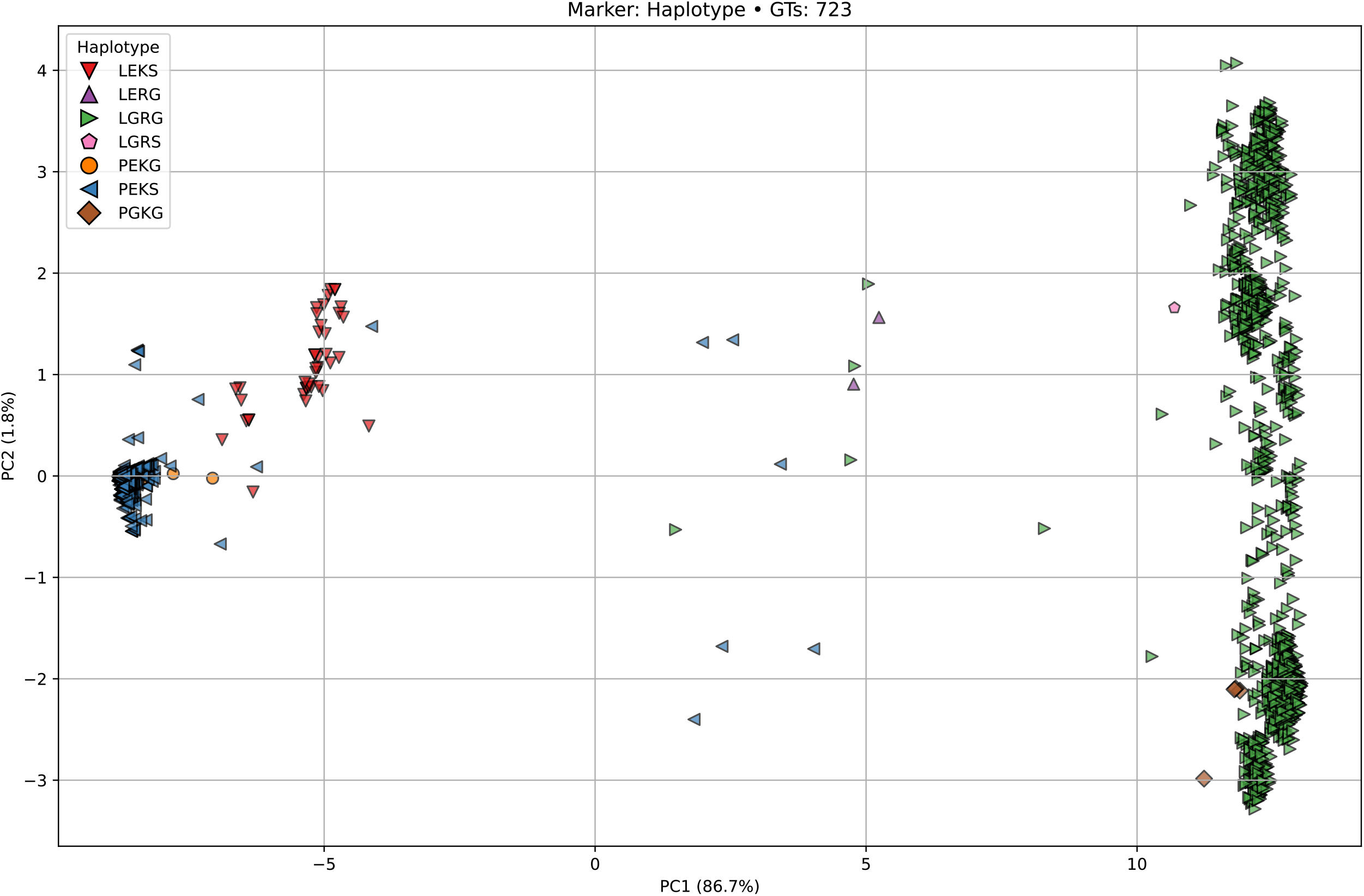
Principal Components Analyses of haplotypes of the *RND2–RNU2* region: (a) African populations (1097 SNPs); (b) European and Asian populations (723 SNPs).

We analysed haplotype frequencies of 2548 individuals available from the 1000 genomes project (Lowy-Gallego et al 2019). LEKS is the majority haplotype in African populations and is also found at low frequency among individuals with South Asian ancestry, with somewhat higher frequency among individuals with ancestry from the south of the sub-continent (Indian Telugu in the UK and Sri Lankan Tamil in the UK) than from the north of the sub-continent (Gujerati Indians in Texas and Punjabis from Pakistan in the UK). LEKS is rare or absent in recent European and East Asian populations. After LEKS, the most common African haplotypes are LGRG, LERG, and PEKS. The frequency of PEKS ranges from 5% in Esan from Nigeria to 12% in Luhya from Kenya. In European and East Asian populations, more than 97% of haplotypes are PEKS or LGRG, with PEKS comprising roughly two-thirds of all haplotypes. In South Asian populations, PEKS and LGRG occur at nearly equal frequencies.

Seven out of seven Neanderthal haplotypes we examined were LEKS but all sequences from 36 ancient humans (30,000–40,000-yrs-old) from Europe and Asia were either PEKS or LGRG, with PEKS the majority haplotype (9 PEKS/PEKS, 6 PEKS/LGRG, 2 LGRG/LGRG diploid genotypes). These observations show that LEKS haplotypes were already present in the ancestral population from which Neanderthals and anatomically modern humans were derived, that neither of the major European and Asian haplotypes (PEKS, LGRG) was derived from Neanderthals, and that the dominance of these haplotypes in recent populations from Europe and Asia was already established 30,000 years ago.

Pairwise genetic distance comparisons (Table 2) show that the ancient PEKS haplotypes are most similar to modern PEKS haplotypes. Similarly, the Neanderthal LEKS-like haplotype is most similar to modern LEKS haplotypes. There is no human or hominin haplotype that has a particularly low genetic distance to the *Pan* (LERS) haplotype. Likely the linkage disequilibrium and human haplotypes originate after the phylogenetic split with *Pan*. Notably LGRG has the largest pairwise genetic distance to the *Pan* haplotype, suggesting increased differentiation due to selection or drift. This is corroborated by a low value of Fay and Wu’s H, representing an excess of high-frequency derived alleles, for LGRG haplotypes across ancestries (Supplementary Table 1).

**Table 2.** Pairwise genetic distances between haplotypes.

|  | PEKS | LEKS | LERG | LGRG | *Pan* | Neanderthal |
| --- | --- | --- | --- | --- | --- | --- |
| LEKS | 0.0156 |  |  |  |  |  |
| LERG | 0.0504 | 0.0485 |  |  |  |  |
| LGRG | 0.0526 | 0.0604 | 0.0389 |  |  |  |
| *Pan* | 0.0664 | 0.0654 | 0.0673 | 0.0772 |  |  |
| Neanderthal | 0.0150 | 0.0123 | 0.0454 | 0.0572 | 0.0618 |  |
| PEKS Ancient | 0.0046 | 0.0171 | 0.0546 | 0.0671 | 0.0682 | 0.0143 |

The databases of modern human sequences we have examined include a small number of haplotypes that are recombinants among the major haplogroups. Although the suppression of recombination does not appear to be absolute, recombination has been sufficiently rare as to have preserved long-range linkage disequilibrium *or* the disruption of favorable epistasis within the region selects against recombinants when they arise *or* some apparent recombinants are errors arising from template-switching during sequencing. The rarity of recombination is corroborated by our finding that SNPs within the region show strong linkage disequilibrium within Africa and near-complete linkage disequilibrium in Eurasia.

PEKS and LGRG haplotypes of *BRCA1* could have been derived from an ancestral LERS haplotype in four mutational steps: (1) Arg1183 (rs16942C) arose by mutation as the progenitor of LEKS haplotypes; (2) Gly1613 (rs1799966C) arose by mutation as the progenitor of LERG haplotypes; (3) Gly1038 (rs16941C) arose by mutation on an LERG haplotype as the progenitor of LGRG haplotypes; and (4) Pro831 (rs799917G) arose by mutation on an LEKS haplotype as the progenitor of PEKS haplotypes. In this schema, the relative order of (1) and (2) is unspecified. The reason for suggesting Pro831 (rs799917G) is a relatively recent mutation is that the derived allele in African populations does not occur on an extended haplotype, with many haplotype-specific SNPs, unlike Gly1038 (rs16941C), Arg1183 (rs16942C), and Gly1613 (rs1799966G).

We conjecture that PC1 in the analyses of Figure 1 captures an ancient balanced polymorphism between LEKS/PEKS and LERG/LGRG haplogroups. In populations outside of Africa, PEKS haplotypes displaced LEKS haplotypes on one side of the polymorphism and LERG haplotypes were lost on the other side. The loss of LERG (a minority African haplotype) could plausibly be explained as a sampling effect arising from an outside-of-Africa bottleneck but the displacement of LEKS (the majority African haplotype) by PEKS (a minority African haplotype) is less easily explained by genetic drift and shows features compatible with a selective sweep, including very low diversity among PEKS haplotypes and a negative value of Tajima’s D (Tajima 1989), representing an excess of rare alleles for PEKS haplotypes (Supplementary Table 1). Recent movements of peoples have generated admixed populations in which European PEKS haplotypes now coexist with African LEKS haplotypes. This raises the question whether the causes of the selective sweep that favored PEKS over LEKS were strictly historical or whether fitness differences between these haplotypes persist in modern admixed populations.

African-American women and women from sub-Saharan Africa exhibit lower overall rates of breast cancer than non-Hispanic White women, but earlier age of onset of aggressive tumors and higher mortality (Fregene and Newman 2005). In a study from Northern California, African–American women with breast cancer had lower rates of *BRCA1* mutations than non-Hispanic white women when averaged across all ages, but a higher rate of *BRCA1* mutations in the subset of women who developed cancer before 35 years (John et al. 2023). Both biological and sociological factors probably play a role in the distinctive epidemiology of breast cancer in women with African ancestry (Dietze et al. 2015). A desideratum for future research is a better understanding of the haplotype structure of the *RND2–RNU2* region within Africa. This gap in current knowledge could be addressed by modern long-read sequencing that will allow determinations of haplotypic phase.

### Genomic structure

Our analyses of genomic structure are based on the CHM13 telomere-to-telomere assembly (Nurk et al. 2022) with ‘proximal’ and ‘distal’ used to specify positions relative to the centromere of human chromosome 17. The region of suppressed recombination contains five protein-coding genes (*CCDC200, TMEM106A, NBR1, BRCA1, RND2*) and is flanked distally by *ARL4D* and proximally by *VAT1* (Figure 2). Synteny from *ARL4D* to *VAT1* has been conserved between humans and chimeras (*Callorhinchus millii*). The most obvious feature of Figure 2 is the regular array of nine *RNU2* repeats between *CCDC200* and *ARL4D*. Also evident is the high density of intronic and intergenic repeats (mainly *Alu* elements) to either side of the *RNU2* locus, with a somewhat higher density on the proximal side.

**Figure 2.**
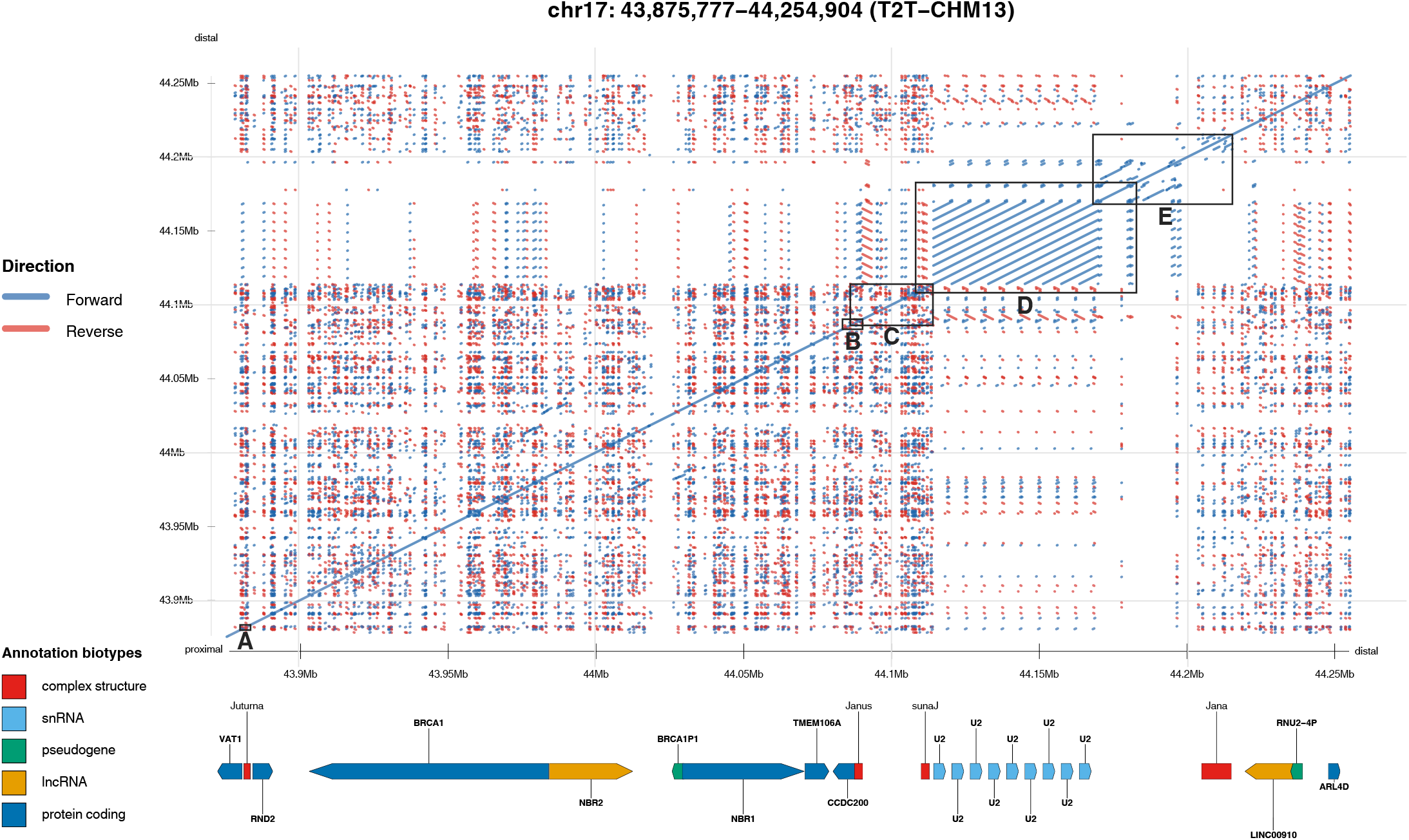
Self-similarity plot of *VAT1* to *ARL4D* showing locations of genes and major structural features. Boxed regions are presented in great detail in other figures: A (*VAT1–RND2* intergenic region) in Figure 7; B (*CCDC200* start site and Janus) in Figure 6; C (proximal flanking region from Janus to sunaJ) in Figure 5; D (*RNU2* array) in Figure 4; E (retroviral-rich region) in Figure 3.

The *RNU2* tandem array was not included in GRCh37 and is inverted in GRCh38 relative to CHM13 with the repeats highly irregular. These differences from CHM13 are likely caused by assembly errors in GRCh38. We found sequences coding for U2 snRNAs between *CCDC200* and *ARL4D* in the genomes of elephants, cows, mice, and primates although there is little detectable similarity of sequences among the major mammalian groups apart from the U2 coding sequences themselves.

Our analysis discovered four multi-kilobase sequences (Jana, Janus, sunaJ, Juturna) in CHM13 that are predicted to form strong secondary structures (Figure 2). In Roman mythology, Janus was the two-faced god of beginnings and endings and Jana and Juturna were mothers of his children. Jana and Juturna form the distal and proximal bookends of the region of suppressed recombination. Janus is centrally placed, gazing proximally past *BRCA1* to Juturna and distally past *RNU2* to Jana, with sunaJ an inverted copy of Janus. Unlike Jana, which is dominated by direct repeats that are conjectured to undergo slipped-strand mispairing, Janus and Juturna contain inverted repeats that are conjectured to form large hairpins of single-stranded DNA, although these structures are barely discernible at the scale of Figure 2.

Slipped-strand structures and single-stranded hairpins create obstacles for DNA replication, especially during lagging-strand synthesis (Khristich and Mirkin 2020). Furthermore, complementarity between DNA sequences entails complementarity between DNA templates and transcribed RNAs. Therefore, Janus, sunaJ and Juturna may also be associated with DNA–RNA hybrids (R-loops) formed during transcription and with conflicts between DNA replication and RNA transcription.

Jana, sunaJ, Janus, and Juturna may be impediments not only to the progression of replication forks at mitotic S phase but also to synapsis of homologues in meiotic prophase. In the absence of synapsis, homologous repair of double-strand breaks is restricted to using sister chromatids as templates. Asynapsis could thus contribute to the suppression of recombination in the genomic region between *ARL4D* and *VAT1*. Two implications of this hypothesis are that BRCA1 protein should be present at the *BRCA1* locus during meiotic prophase and that asynapsis of the *RND2–RNU2* supergene is tolerated in both male and female meiosis.

### Between *ARL4D* and *CCDC200* (from Jana to Janus)

The genomic region between *ARL4D* and *CCDC200* will be divided into four subregions: a ‘distal flanking region’ between *ARL4D* and Jana; a ‘retroviral-rich region’ from Jana to the distal end of the *RNU2* array; the ‘*RNU2* array’; and a ‘proximal flanking region’ from the proximal end of the *RNU2* array to Janus and the translation start site of *CCDC200* (see Figure 2). The distal and proximal flanking region are characterized by a high density of repeats (mainly *Alu* elements) with a somewhat higher density of repeats in the proximal flanking region. The retroviral-rich region extends from Jana to the beginning of the array of nine *RNU2* repeats. The major direct repeats observable immediately adjacent to the *RNU2* array are two endogenous retroviruses (ERVs). The retroviral-rich region and *RNU2* array are dominated by direct repeats. The presence of many direct repeats without inverted repeats suggests that this region of the genome may be subject to slipped-strand mispairing. This contrasts with the distal and proximal flanking regions that contain a high density of both direct and indirect repeats. Thus, a central ‘slippery’ region is flanked by ‘sticky’ sequences. Subsections (i)–(iv) below describe the four subregions of CHM13. Subsection (v) offers a general overview and comparison of CHM13 and GRCh38.

#### (i) Distal flanking region

*ARL4D* lies outside the region of high linkage disequilibrium and suppressed recombination. In CEU families from Utah, rs12946015 from the 3’ UTR of *ARL4D* is in weak linkage disequilibrium with rs799917 from exon 11 of *BRCA1* (R^2^ = 0.05). Proximal to *ARL4D* is a 34-kb region of sequences derived from degenerating *RNU2* repeats that have been disrupted by insertions of numerous transposable elements. These *RNU2*-related sequences are mostly inverted relative to the tandem array. The distal flanking region consists of 61% interspersed repeats (33% SINEs, 11% LINEs, 14% ERVs) including 40 *Alu* insertions in more or less random orientation with respect to each other. The *RNU2*-related sequences include *RNU2-4P*, a U2 snRNA pseudogene, and a long terminal repeat (LTR) that has been disrupted by an *Alu* insertion. Long-noncoding RNAs transcribed from this region are known as *LINC00910*. The distal flanking region is well conserved in the genomes of great apes. There is 99% identity of the human sequence to bonobo and chimpanzee sequences; 98% to gorilla; and 96% to orangutan. SNPs from this region (e.g., rs34127860, rs12600401) are in weak linkage disequilibrium with SNPs in *BRCA1*.

#### (ii) Retroviral-rich xsregion

Proximal to the distal flanking region is a 45-kb region dominated by retroviral sequences (78% ERVs; 6% SINEs) that is notable for an absence of inverted repeats and a paucity of *Alu* elements (Figure 3). The dearth of *Alu* elements suggests either that retroviral integrations into this region occurred more recently than the insertions of *Alu* elements in the *RNU2*-related sequences of the proximal flanking region or that *Alu* insertions are not tolerated because they disrupt important functional elements.

**Figure 3.**
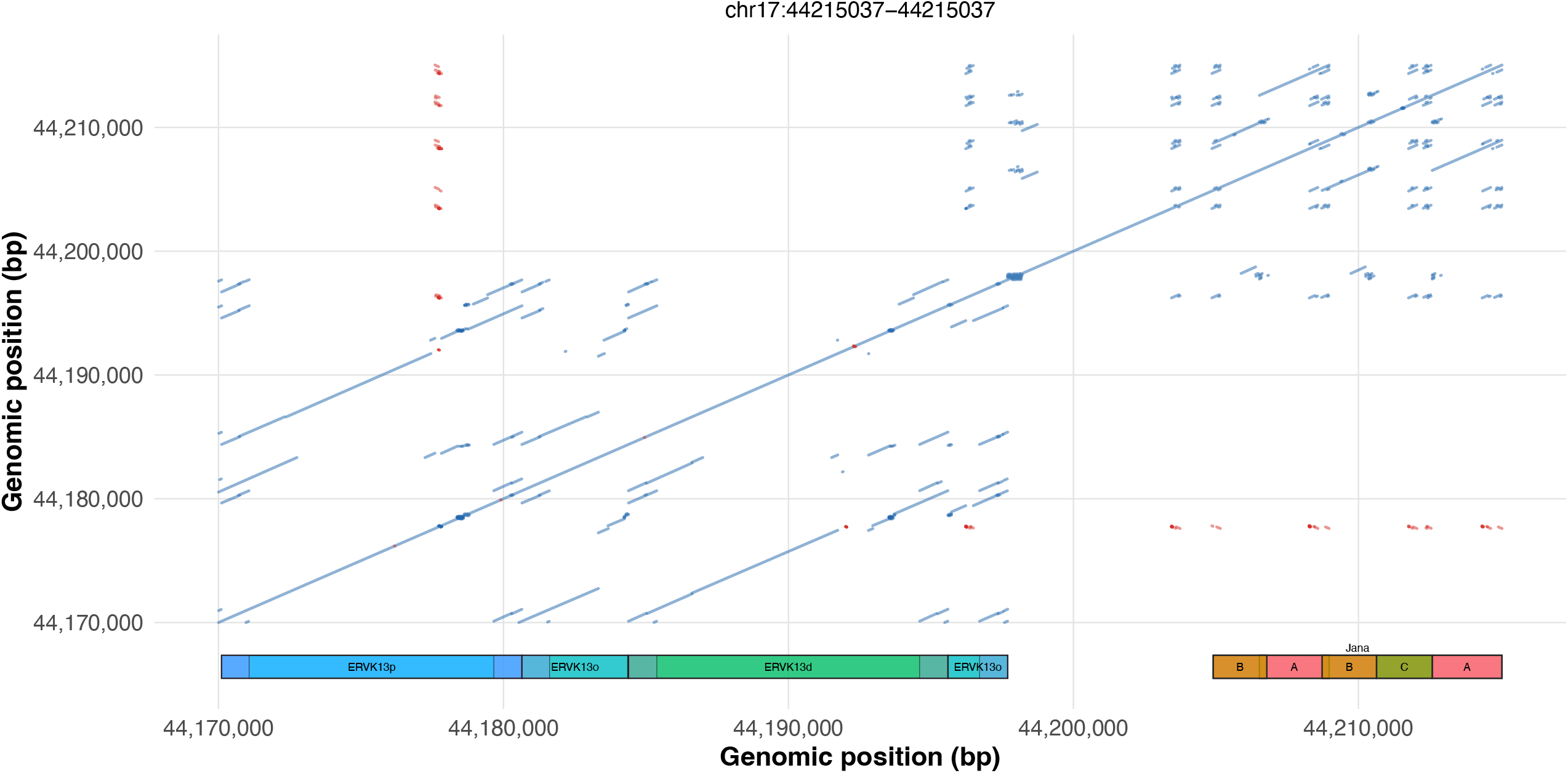
Self-similarity plot of the retroviral-rich region with the long version of Jana (upper right). Three ERVK13 retroviruses (lower left) comprise the proximal two-thirds of this region.

Jana, which is located in the distal third of the retroviral-rich region, is polymorphic for a structural variant. The long variant of Jana (present in CHM13) is 10.1 kb in length and consists of two copies of a 2.4-kb ‘A sequence’, two copies of a 1.7-kb ‘B sequence’, and a single-copy of a 2.6-kb ‘C sequence’, with one copy of A inserted between the two copies of B in the order ACBAB. Each A sequence contains two *Alu* elements (24%) and sequences derived from an ERVL element (13%). Each B sequence contains an *Alu* element (13%) and sequences derived from a PRIMA41 ERV and MER41C LTR (60%). The C sequence contains two *Alu* elements (32%). The short variant (present in GRCh38) is 6.3 kb in length and consists of single copies of A, B, and C in the order ACB. Details of the repeat composition of the structural variants and our attempt to unravel their complex history are presented in Box 1. A third MER41C LTR is located 6 kb from the proximal B sequence, 500 bp distal to the first of six LTRs associated with ERVK13 retroviruses (see below).

All PEKS haplotypes possess the short variant. Most LEKS and LGRG haplotypes possess the long variant although a few possess the short variant, providing evidence of between *BRCA1* and Jana. In European and East Asian populations, SNPs immediately proximal to Jana (e.g., rs4793234, rs4793233) are in strong LD with SNPs tagging the PEKS/LGRG dimorphism (e.g., rs799917, rs16942). Thus, Jana appears to form the distal boundary of the region of maximal linkage equilibrium. We conjecture that the long variant is susceptible to slipped-strand mispairing which creates particular difficulties for DNA replication and meiotic synapsis.

The proximal two-thirds (29-kb) of the retroviral-rich region contains three ERVK13 retroviruses associated with six LTRs of 72–91% identity. The six LTRs are in the same orientation as the LTRs of the *RNU2* repeats in CHM13 but are in inverted orientation in GRCh38. All these LTRs belong to the LTR13 family of Liao et al. (1998). Two of the retroviruses are 10.6-kb in length, possess LTRs at both ends, and are 87% identical to each other. We will name them the distal and proximal ERVs (ERVK13d and ERVK13p). The flanking LTRs of ERVK13d are 91% identical to each other whereas the flanking LTRs of ERVK13p are 73% identical to each other. ERVK13d has integrated into an older retrovirus (ERVK13o) at a sequence predicted to form slipped-strand structures in ERVK13o, ERVK13d, and ERVK13p. The proximal LTR of ERVK13o is immediately adjacent to the distal LTR of ERVK13p and the proximal LTR of ERVK13p immediately adjacent to the first LTR of the *RNU2* array. Thus, ERVK13p is flanked on both sides by tandem pairs of LTRs. These retroviruses are discussed in Box 3.

#### (iii) *RNU2* array

Figure 4 shows the *RNU2* array of CHM13 in its immediate genomic context. Although this array contains nine copies of the 6.1-kb monomer, the number of *RNU2* repeats within an array is highly variable with from 5 to 82 6.1-kb repeats in a sample of 41 unrelated human chromosomes (Tessereau et al. 2014). When array sizes were compared between distantly-related individuals with the same *BRCA1* founder mutation, changes in copy number occurred about once every 200 generations, with these changes involving multiple repeats, for example a change in number of 10 repeats (Tessereau et al. 2014).

**Figure 4.**
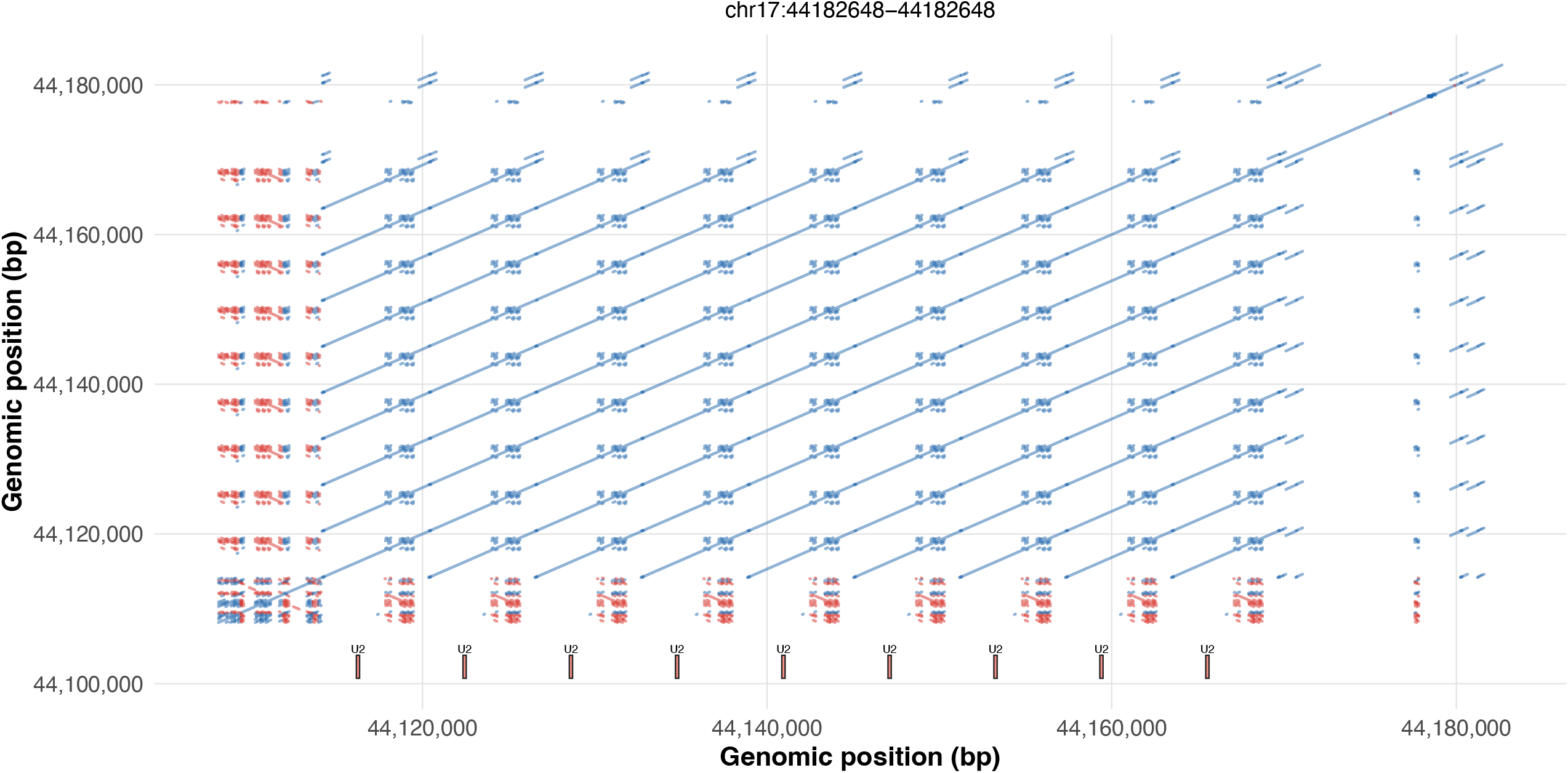
Self-similarity plot of the *RNU2* array located between sunaJ (lower left) and ERVK13p (upper right). The regularly spaced LTR repeats can be seen across the top of the plot aligning with the doubled LTRs flanking ERVK13p. A different segment of the *RNU2* repeat, containing three *Alu* elements, is shared with sunaJ. This segment can be visualized across the bottom of the plot.

The arrays of *RNU2* repeats are highly regular in CHM13 and the bonobo reference genome, suggesting ongoing concerted evolution. CHM13’s array contains three variants of the 188-bp U2 snRNA. Proceeding from the proximal end of the array, there are four copies of canonical *RNU2-1* followed by three copies with an A-to-G substitution at nucleotide 160, and then two copies with a C-to-T substitution at nucleotide 18 in addition to the A-to-G substitution at nucleotide 160. Whatever the mechanism of concerted evolution, it appears to have involved contiguous expansions of variants within the array.

Each of the nine 6.1-kb repeats possesses an ERVK13 LTR, a 188-bp U2 snRNA coding sequence, and a polypyrimidine microsatellite. The polypyrimidine microsatellite contains mirror symmetries and is predicted to form ‘H-DNA’ triplexes in which part of the polypyrimidine strand folds into the major groove of the DNA duplex leaving the complementary polypurine sequence single-stranded (Hisey et al. 2024). Between successive LTRs, are 5.1 kb of sequence containing 67% interspersed repeats recognized by RepeatMasker, including sequences derived from an ERVL retrovirus, L1 and L2 LINEs, *Alu* and *MIR* SINEs, and DNA transposons. The four *Alu* elements are all in the same orientation. The remaining 33% of the inter-LTR sequence contains *RNU2* ‘specific’ sequences including the U2 coding sequence and polypyrimidine microsatellite.

Human and great ape reference sequences contain tandem arrays of 6.1-kb *RNU2* repeats. Old World monkeys are reported to possess an 11-kb *RNU2* repeat (Matera et al. 1990). We found single copies of this 11-kb unit in reference sequences of Old World monkeys but not tandem arrays. This may be an assembly problem analogous to the absence of the *RNU2* array in GRCh37. In baboons, each copy of U2 snDNA was reported to be associated with part of an ERVK13 retrovirus flanked by two LTRs (Pavelitz et al. 1995). These retroviral sequences are similar to the human sequence we call ERVK13o. The 6-kb repeat unit of apes could be derived from an 11-kb repeat similar to that reported for Old World monkeys by recombination between the two LTRs of this retrovirus to produce a solitary LTR associated with a U2 coding sequence (Pavelitz et al. 1995). U2 coding sequences of platyrrhine primates (New World monkeys) and mice are not associated with LTRs nor with tandem arrays. Therefore, these appear to be derived features of the *RNU2* loci of catarrhine primates (Old World monkeys and apes).

One possible mechanism of concerted evolution would involve unequal recombination within the array resulting either from slipped-strand mispairing between complementary strands of a single chromatid or unequal ‘crossing over’ between sister chromatids facilitated by the general ‘slipperiness’ of the *RNU2* array. Recombination within an array could also generate extrachromosomal circles containing one or more repeats that could be reincorporated into arrays by homologous recombination. Another possible mechanism would involve targeted retrotransposition into the array. In this mechanism, one or more *RNU2* repeats flanked by two LTRs would behave as a non-autonomous retrotransposon. An analogy is provided by the R2 retrotransposons that play a role in the concerted evolution of *Drosophila* rDNA repeats (Nelson et al. 2023).

The association of U2 genes with retroviral LTRs originated before the divergence of Old World monkeys from humans and apes as evidenced by the conservation of LTRs associated with *RNU2* repeats in both groups. The LTRs of human *RNU2* repeats exhibit 86% sequence identity to the LTRs of baboon *RNU2* repeats and can be inferred to have retained a conserved function. Another feature conserved between humans and baboons is the presence of the ERVK13p provirus in close proximity to the repeats.

#### (iv) Proximal flanking region

The proximal flanking region has a 4-kb inverted duplication at its proximal and distal ends (Figure 5). This duplication contains the sequence we call Janus, the first coding exon of *CCDC200*, and 1-kb of adjacent intron. The proximal copy of Janus, adjacent to *CCDC200*, will retain the name Janus and we will use sunaJ to designate the inverted copy, adjacent to the *RNU2* array. The 4-kb duplication is absent from the reference sequences of bonobos and baboons, and the region between Janus and sunaJ is inverted in CHM13 relative to the bonobo reference sequence.

**Figure 5.**
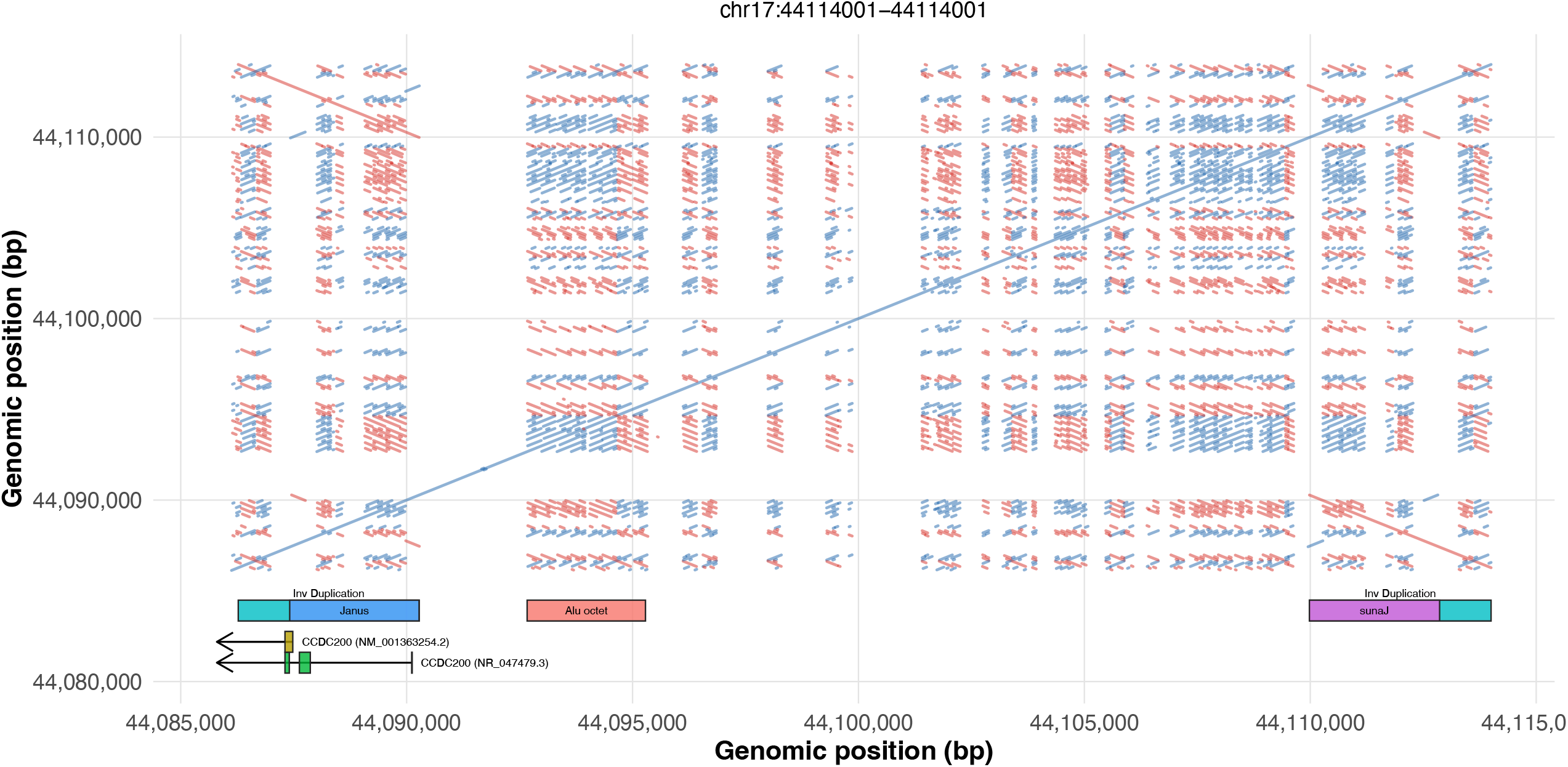
Self-similarity plot of the proximal flanking region showing a 4-kb inverted duplication. One copy of this duplication (lower left) contains Janus and the beginning of *CCDC200* (see Figure 6 for greater detail). The other copy of the duplication (upper right) contains sunaJ. Between the duplicated sequences are multiple *Alu* elements including the *Alu* octet. Potential structure-forming features and *CCDC200* exons are annotated.

Our description of the proximal flanking region will begin with the simpler arrangement of the bonobo reference sequence in which there are 24 kb (55% SINEs, 2% LINEs, 13% ERVs) between Janus and the most proximal LTR of the *RNU2* array. An octet of eight *Alu* elements (seven *AluSx* and one *AluY*) begins barely 300 bp from the end of the *RNU2* array, six aligned in direct orientation followed by two in inverse orientation. The *Alu* octet thus marks a transition from the ‘slippery’ retroviral-rich region and *RNU2* array, in which almost all repetitive sequences are in direct orientation, to a ‘sticky’ region of the genome in which inverted repeats are predicted to form hairpins of single-stranded DNA. (The baboon sequence has five *AluSx* elements, four in direct orientation followed by one in inverse orientation.) Also notable is a ‘super slippery’ array of 11 *AluSx* elements, all in the same orientation, 2 kb distal to Janus. (The corresponding region of the human sequence contains six *AluSx* elements).

Janus is a 2.8-kb sequence that encompasses the 5’ untranslated region and beginning of the first coding exon of *CCDC200*. It is bookended by non-*Alu* related inverted repeats that are complementary for 296 out of 301 nucleotides and are predicted to form the stems of a large hairpin. We will call these the distal and proximal faces of Janus. Between the two faces of Janus are 2.2 kilobases containing six *Alu* elements (five direct repeats followed by an element in inverse orientation). Four of the *Alu* elements, including the element in inverse orientation, belong to the *AluSx* family. A kilobase of this sequence, including three of the *Alu* direct repeats, are highly similar to part of an *RNU2* repeat that contains neither the U2 snRNA coding sequence nor retroviral LTR. The region between the two faces of Janus has the potential to form complex structures including slipped-strand mispairing between the direct *Alu* repeats and hairpins formed by pairing of the inverted *Alu* element with the direct repeats (hairpins within the Janus hairpin). The evolutionary history of Janus is discussed in Box 2.

Figure 6 shows a self-similarity plot of the region surrounding the *CCDC200* start site with Janus to the upper right and the adjacent intron to the lower left. The intron contains a ‘tangle’ of nine *Alu* elements with three pairs of inverted repeats. The first coding exon of *CCDC200* is thus embedded within a snarl of complementary sequences that are predicted to form strong secondary structures of single-stranded DNA and DNA–RNA hybrids (R-loops).

**Figure 6.**
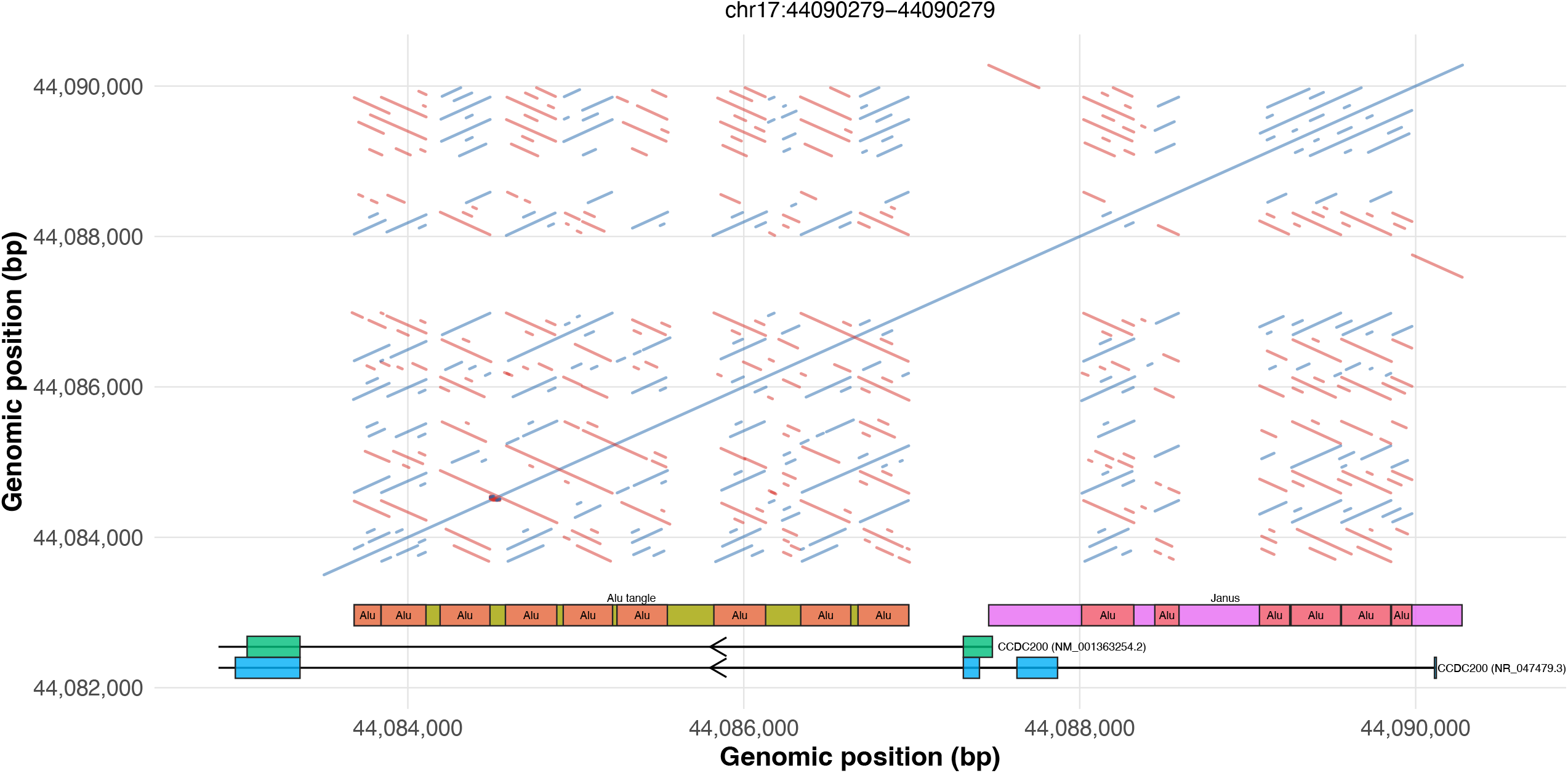
The region surrounding the translation start-site of *CCDC200* with Janus at upper right and the *Alu* tangle at lower left.

The *RNU2*-related sequences contained within the human and baboon Janus hairpins exhibit 88% identity to each other whereas the human *RNU2*-related sequence shows 92% identity to a human *RNU2* repeat and 87% identity to a baboon *RNU2* repeat. Although these comparisons are based on a small number of sites, they suggest the possibility of a functional interaction, based on sequence complementarity, between the Janus hairpin and the *RNU2* repeats, as well as an exchange of information between Janus and the *RNU2* repeats since apes diverged from Old World monkeys.

Transcription of human *CCDC200* proceeds proximally from within Janus toward *TMEM106A*. The proximal face of Janus is only 45 nucleotides from the AUG codon that initiates translation of CCDC200 protein. Genbank contains two transcripts of *CCDC200* (NM_001363254, NR_047479) that differ because of alternative splicing within Janus. NR_047479 has two upstream exons that are spliced into the first exon of NM_001363254 after the latter’s AUG start codon. NR_047479 is labelled a long noncoding RNA, but the twelfth codon of NM_001363254 is another AUG that may provide a start codon for the NR_047479 transcript. Non-coding transcripts that include parts of Janus have been described as *LINC00854* and *TMEM106A-AS1*.

The proximal breakpoint of the 4-kb duplication shown in Figure 5 occurs within the *Alu* tangle and the distal breakpoint occurs at the distal end of Janus. A copy of the 4-kb duplication is included within a 34-kb inversion by which CHM13 differs from the bonobo reference sequence. One possible scenario is that a tandem duplication of 4-kb in a human ancestor was followed by a 34-kb inversion with its proximal breakpoint between the duplicated sequences. As a result of this complex rearrangement (or rearrangements), a copy of Janus is present just outside the proximal end of the inversion and an inverted copy of Janus (sunaJ) is present just inside the distal end of the inversion.

From its distal end, the 34-kb inversion is comprised of the inverted copy of the 4-kb duplication, sequences orthologous to the bonobo distal flanking region including the *Alu* octet (28 kb) and 2 kb of an *RNU2* repeat, including a truncated LTR that is located 1 kb from the proximal end of the inversion. Because of this inversion, the proximal flanking region has been extended in CHM13, relative to the bonobo proximal flanking region, by the 4-kb duplication and incorporation of 2-kb sequences that were once part of the regular *RNU2* array but are now separated from, and inverted relative to, that array. One consequence is that sunaJ, rather than the *Alu* octet, is immediately adjacent to the proximal end of the *RNU2* array.

#### (v) Synopsis of region between *ARL4D* and *CCDC200*

The distal and proximal flanking regions of CHM13 contain *RNU2*-related sequences, interspersed with numerous *Alu* elements in a mixture of inverted and direct orientations. In contrast, the retroviral-rich region and *RNU2* array are characterized by a paucity of *Alu* elements, absence of inverted repeats, and abundance of directly repeated sequences at scales from a few nucleotides to multiple kilobases. Because of the dominance of direct repeats, we conjecture that the two strands of DNA are predisposed to slip past each other into out-of-register pairings.

*RNU2* repeats within the tandem array of CHM13 are almost identical to each other, suggesting that the arrays have undergone processes of homogenization (concerted evolution). *RNU2*-related sequences in the flanking regions are mostly in inverted orientation relative to the *RNU2* array, have diverged from the canonical repeats, and do not encode functional U2 snRNAs. We conjecture two ongoing processes disrupt concerted evolution of *RNU2* repeats. The first is a predisposition for endogenous retroviruses to insert into the *RNU2* array. The second are inversions that remove *RNU2*-related sequences from engagement in whatever processes are responsible for their concerted evolution. The general organization resembles that of centromeres where there is concerted evolution of repeats at centromeric cores with degeneration of repeats in pericentromeric flanks (Hartley and O’Neill 2019; Miga and Alexandrov 2021; Haig 2022).

The distal flanking regions and 36 kilobases of retroviral-rich sequence proximal to Jana are 99% identical in CHM13 and GRCh38. Sandwiched between these regions of high identity, CHM13 possesses the long variant of Jana (10.1 kb) and GRCh38 possesses the short variant of Jana (6.3 kb). Proximal to the retroviral-rich region, GRCh38 differs from CHM13 by an inversion of the region between ERVK13p and the *CCDC200* start site that includes the entire *RNU2* locus. Whereas the *RNU2* array is highly regular in CHM13 it is irregular in GRCh38. The GRCh38-specific inversion and irregularity of the *RNU2* array are probably assembly errors of this highly repetitive part of the genome.

### Between *CCDC200* and *VAT1* (from Janus to Juturna)

The most notable feature of the 207-kb region that lies between *CCDC200* and *VAT1* is the high density of repetitive elements (see Figure 2). This section summarizes the region’s major features.

*CCDC200* abuts *TMEM106A* tail-to-tail with the 3’ ends of *CCDC200* and *TMEM106A* separated by 1.3 kb. *TMEM106A* abuts *NBR1* head-to-tail with the 5’ end of *TMEM106A* separated by 2.6 kb from the 3’ end of *NBR1*. In most mammals, *NBR1* and *BRCA1* share a bidirectional promoter. This promoter has been tandemly duplicated in catarrhine primates (Barker et al. 1996; Brown et al. 1996). The distal duplicate functions as a bidirectional promoter for *NBR1* and *BRCA1P1* (a pseudogene highly similar to the 5’ end of *BRCA1*) whereas the proximal duplicate functions as a bidirectional promoter for *BRCA1* and *NBR2* (a long noncoding RNA whose 5’ end is highly similar to *NBR1*).

The initial duplication of the bidirectional promoter and surrounding sequences was about 4.4 kb, with the distal end of the distal duplicate located in the third intron of *NBR1* and the proximal end of the proximal duplicate located in the second intron of *BRCA1*. Since this initial duplication, a 7.4-kb Harlequin endogenous retrovirus has inserted into the distal duplicate. Between the proximal end of the distal duplicate and the distal end of the proximal duplicate resides 32 kb of sequence much of which aligns to *PLEKHM1* and *LRRC37A4P* from one flank of the *MAPT* inversion at 17q21.31 and to intronic sequences from *LRRC37A* on the other flank of the *MAPT* inversion (Stefansson et al. 2005). *LRRC37A*-related genes are present at several locations on human chromosome 17 as parts of larger segmental duplications (Jin et al. 2004; Zody et al. 2008; Giannuzzi et al. 2013). Recombination between these sequences is a cause of recurrent rearrangements of chromosome 17 (Cruts et al. 2005; Kehrer-Sawatzki et al. 2017). The presence of *PLEKHM1* and *LRRC37A4P*-related sequences between the bidirectional promoters of *BRCA1/NBR2* and *BRCA1P1/NBR1* suggests that copies of the *LRRC37A*-segmental duplication were independently inserted into the second intron of *BRCA1* and third intron of *NBR1* with pairing and recombination between them responsible for the *BRCA1/NBR1* duplication.

Human *BRCA1* is notable for abundant *Alu* elements that comprise more than 40% of the gene’s sequence (Smith et al. 1996). Another 5% of the *BRCA1* sequence is derived from LINE1 and LINE2 elements. Closely-spaced *Alu*s are predicted to cause slipped-strand mispairing when *Alu*s occur in the same orientation and hairpins when *Alu*s occur in inverted orientation. These secondary structures may be sources of replicative stress. Abundant intronic *Alu* elements probably also create difficulties in splicing because *Alu* elements contain cryptic splice sites (Zarnack et al. 2013; Haig 2025).

*RND2* lies immediately proximal to the 3’ end of *BRCA1* in a head-to-head arrangement with *VAT1*. RND2 lies just within, and VAT1 lies just outside, the proximal boundary of the region of maximal linkage disequilibrium.

SNPs within *RND2* are in strong LD with distant SNPs on the far side of *RNU2* but more weakly associated with much closer SNPs within *VAT1*. In CHM13, the human *VAT1*–*RND2* intergenic region is 3.5-kb and contains a cluster of seven *Alu* elements that exhibit three reversals of orientation between adjacent elements (Figure 7). This 2.2-kb cluster of *Alu* elements is the sequence we call Juturna. It is predicted to form strong single-stranded hairpins within hairpins. In GRCh38, the *VAT1–RND2* intergenic region is only 2.8 kb because GRCh38 lacks a pair of inverted *Alu* elements present in CHM13 that are predicted to form the strongest internal hairpin of Juturna. We believe that absence of this internal hairpin is an assembly error, arising from difficulties in sequencing, rather than a polymorphism in the human population. All seven *Alu* elements are present in the *VAT1–RND2* intergenic regions of great apes. Multiple *Alu* elements that are predicted to form strong secondary structures are also present in the *VAT1*–*RND2* intergenic regions of gibbons, Old World monkeys and New World monkeys. In marked contrast, the *VAT1*–*RND2* intergenic regions of lemurs (*Microcebus*) and mice (*Mus*) do not contain repeats.

**Figure 7.**
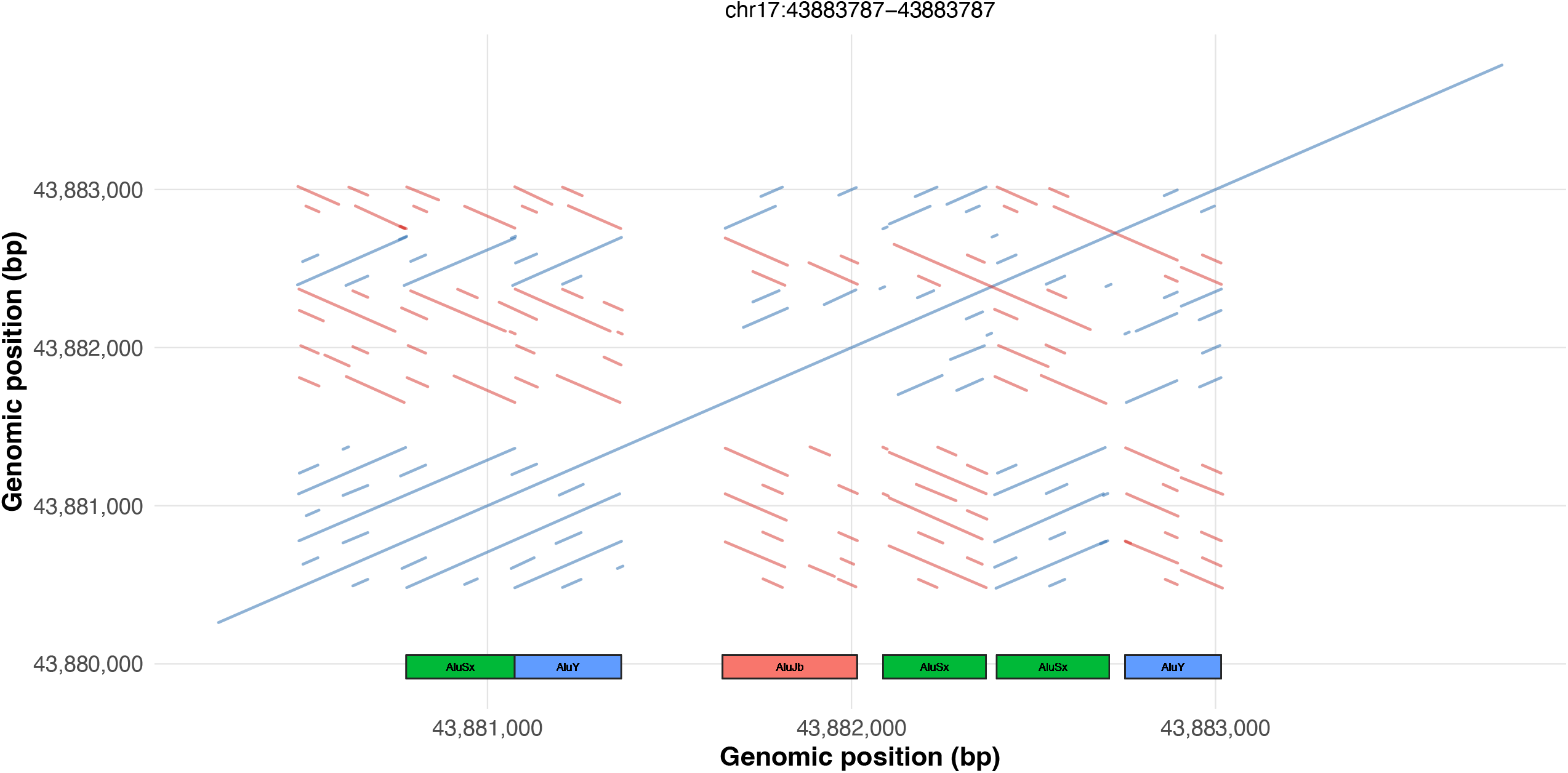
Self-similarity plot of the *VAT1*–*RND2* intergenic region containing Juturna. *Alu* elements are annotated.

### Selection in a “supergene”

Supergenes have been defined as genomic regions containing sets of tightly linked loci that control phenotypic polymorphisms maintained by balancing selection (Gutiérrez-Valencia et al. 2021). The *RND2–RNU2* region satisfies these criteria: its genes are tightly linked and the presence of two major haplogroups in all modern human populations suggests that the polymorphism is maintained by some form of balancing selection. *RND2–RNU2* thus joins a growing list of human ‘supergenes’ that includes the major histocompatibility complex (Kulski et al. 2022), the killer cell inhibitory superlocus (Gourraud et al. 2010), and the *MAPT* inversion (Campoy et al. 2022).

If the *RND2–RNU2* region is subject to balancing selection, then this implies that its major haplogroups are functionally non-equivalent. The Ensembl Variant Effect Predictor (McLaren et al. 2016) classifies five out of the 365 SNPs that differ between the LGRG and PEKS consensus haplotypes as missense variants, four of which occur in *BRCA1* (the tagging SNPs of Table 1) and one in *NBR1*. Functional predictions by Polyphen (Adzhubei et al. 2010) classify four of the five as benign and one as possibly damaging. Genome-wide association studies of haplogroup-specific SNPs reveal that PEKS, when compared to LGRG, is associated with earlier menopause (Day et al. 2015, Ruth et al. 2021) and earlier menarche (Horikoshi et al. 2018), higher body mass index (Hawkes et al. 2023; Huang et al. 2022) and longer telomeres (Burren et al. 2024). *BRCA1* of PEKS haplotypes is transcribed at higher levels than *BRCA1* of LGRG haplotypes (Cox et al. 2011). Moreover, in 97 out of 104 eQTL hits to the region, expression was higher from PEKS haplotypes than from LGRG haplotypes and splicing QTL studies also found tissue-dependent alternative splicing between the PEKS and LGRG haplotypes, primarily involving *NBR1* and *NBR2* transcripts (GTEx Consortium 2020). What is currently lacking is an understanding of the phenotypes under selection.

A necessary condition for a balanced dimorphism is that each ‘allele’ has higher genic fitness when rare than the other ‘allele’ has when common. The simplest way to satisfy this condition is for each ‘allele’ to have higher genic fitness in heterozygotes than the other ‘allele’ has in homozygotes. Genic, rather than genotypic, fitnesses are specified because the two alleles of heterozygotes could have different genic fitnesses as occurs, for example, in segregation distortion. At selective equilibrium, the two alleles would have equal genic fitnesses averaged across all diploid genotypes. In European and East Asian populations, PEKS homozygotes are more common than LGRG homozygotes which implies the former have higher genotypic fitness than the latter. This may explain Osorio et al.’s (2003) observation that pathogenic germline mutations of *BRCA1* are overrepresented on LGRG haplotypes (haplotype II in their paper). Our rationale for this suggestion is that selection against new deleterious recessive mutations is attenuated on LGRG haplotypes because these chromosomes already carry a higher frequency of variants that are deleterious when homozygous. In other words, positive epistasis exists among deleterious recessive mutations under the principle that individuals can only die once.

Epistatic interactions among closely-linked genes favor suppression of recombination to preserve synergistic allelic combinations, and epistasis within haplotypes accumulates after recombination is suppressed because new mutations are restricted to a single genetic background. Epistasis for fitness and suppression of recombination are thus self-reinforcing. In regions of low recombination, positive epistasis is generated between deleterious recessive mutations resulting in ‘pseudo-overdominance’ of complementary haplotypes (Sianta et al. 2023). An extreme endpoint of this dynamic would be a balanced-lethal system of obligate heterozygosity (Wielstra 2020). Because recombination is suppressed across the *RND2–RNU2* supergene, different haplotypes will be associated with different mutations. Whatever the original source of balancing selection that maintained two haplogroups, subsequent accumulation of deleterious recessives will have generated pseudo-overdominance, reinforcing the polymorphism (Wielstra 2020). Because non-recombining haplotypes are inherited as if they were alleles at a single locus, epistatic interactions within haplotypes contribute to additive fitness differences in Mendelian inheritance.

*BRCA1* has meiotic functions that may be subject to cellular selection in both male and female germ lines. Among women undergoing assisted reproduction, women with pathogenic *BRCA1* mutations possess fewer ovarian follicles than age-matched controls (Lin et al. 2017; Lambertini et al. 2018; Helbling-Leclerc et al. 2024) and SNPs in *BRCA1* are correlated with self-reported age at natural menopause in women of European descent (Day et al. 2015). Earlier menopause is thought to be associated with more rapid depletion of the ovarian reserve of oocytes. The variant associated with earlier menopause (rs1799949G) resides on PEKS haplotypes, suggesting that oocytes of women with PEKS haplotypes experience higher rates of atresia than women without PEKS haplotypes.

Spermatocytes with *Brca1* loss-of-function fail to progress beyond pachytene in male mice (Cressman et al. 1999; Xu et al. 2003). At early leptotene in male and female mice, BRCA1 is essential for loading meiotic recombinases at sites of future crossovers (Bai et al. 2024; Jiao et al. 2025). Moreoever, BRCA1 is lost from chromosomal segments after they synapse at zygotene but accumulates on segments that fail to synapse (Mahadevaiah et al. 2008; Broering et al. 2014; Alavattam et al. 2022; Bai et al. 2024). At the latter locations, BRCA1 initiates meiotic silencing of unsynapsed chromosomes (MSUC). Failure of MSUC initiates apoptosis with BRCA1 appearing to be a limiting factor (Mahadevaiah et al. 2008; Kouznetsova et al. 2009). These observations suggest a model in which meiocytes tolerate unsynapsed chromatin silenced by BRCA1 but undergo apoptosis if unsynapsed chromosome segments exceed the limited supply of BRCA1. The present paper argues that human *BRCA1* may reside in an unsynapsed region of the genome and therefore may itself be subject to MSUC.

*BRCA1* has functions in mitotic DNA replication. BRCA1 facilitates replication of gaps of single-stranded DNA and protects stalled replication forks (Nagaraju and Scully 2007; Daza-Martin et al. 2019; Panzarino et al. 2021; Thakar and Moldovan 2021; Schreuder et al. 2024) with the quantity of BRCA1 a limiting factor in the performance of these tasks. Moreover, *BRCA1* is haploinsufficient under conditions of replicative stress associated with stalled replication forks (Pathania et al. 2014, Li et al. 2024). Replication and transcription interfere with each other when they compete for the same DNA template, with close encounters between replisomes and RNA polymerases a cause of replicative stress and double-strand breaks (Browning and Merrikh 2024).

One aspect of the supergene’s phenotype subject to selection may be its own replication. *BRCA1* and *RNU2* play important roles in each other’s replication and each other’s transcription. Moreoever, the *RND2–RNU2* region is replete with sequences that challenge specific functions of BRCA1 protein and U2 snRNA. The wild-type allele of *BRCA1* undergoes frequent somatic mutation in cells heterozygous for a mutated copy of *BRCA1* because of failures of homologous recombination and stalling of replication forks within the *BRCA1* gene (Deshpande et al. 2022). For this reason, *BRCA1* has been said to protect against its own fragility (Martin and McVey 2022). *BRCA1* also protects against metaphase fragility of persistently single-stranded template DNA at the hypertranscribed *RNU2* locus (Pavelitz et al. 2008). Transgenic *RNU2* arrays induce chromosome fragility at ectopic sites (Li et al. 1993).

In addition to the *RNU2* tandem array, the human genome contains *RNU2-2*, present as a single functional gene copy on chromosome 11. Mutations in *RNU2-2* were recently shown to be associated with a dominant neurodevelopmental disorder (Greene et al. 2025; Jackson et al. 2025), with the single-copy *RNU2-2* gene expressed at higher levels in developing brain and retina than were the multiple copies of *RNU2-1* from the *RNU2* tandem array (Jackson et al. 2025). Even more recently, mutations in *RNU2–2* were shown to be associated with a recessive neurodevelopmental disorder (Greene et al. 2025a). Recessive inheritance in these patients shows that the absence of functional U2 snRNA encoded by the *RNU2–1* genes of the *RNU2* tandem array does not compensate for the absence of a functional copy of *RNU2–2*. Perhaps there is a division of labor between *RNU2–1* and *RNU2–2* in different cell types. Two alleles of *RNU2–2* appear sufficient to produce enough U2 snRNA to meet cellular demands in the developing brain. Moreover, the number of *RNU2–1* gene copies in the *RNU2* tandem array is highly variable (Tessereau et al. 2013, 2014). Why, then, do some individuals possess so many functional copies of *RNU2–1*?

R-loops form when a nascent transcript anneals with the DNA template from which it has been transcribed creating a DNA–RNA hybrid duplex and leaving the non-template strand as single-stranded DNA. Such structures create problems for efficient transcription and DNA replication (Crossley et al. 2019). Among other functions, BRCA1 participates in the processing of R-loops at stalled replication forks (Hatchi et al. 2015; Chakraborty and Hiom 2021; Patel et al. 2023). R-loops may create particular difficulties for replication of the *RND2–RNU2* supergene because of the presence of complex secondary structures associated with Jana, Janus and Juturna, with the array of *RNU2* repeats, and with the high density of intronic *Alu* elements within *BRCA1*.

Haig (2025) proposed that genes with roles in DNA replication and repair, including *BRCA1*, evolve features that create difficulties for DNA replication and repair. Replicative stress then tests the competence of a locus’s own gene products to pass a self-imposed examination. Haig’s (2025) rationale for the ‘stress-test hypothesis’ was that the sequence of a gene whose protein product performs a function with high competence has never been exposed to the absence of proteins of high competence for that function. Therefore, features that challenge the competence of a protein are preferentially located in the vicinity of elite alleles because similar features elsewhere in the genome are eliminated by germline selection when they occur in individuals or cells possessing two alleles of lesser competence.

Because BRCA1 facilitates replication of gaps of single-stranded DNA, protects stalled replication forks, and processes R-loops, the stress-test hypothesis predicts that the *RND2–RNU2* supergene will be associated with sequences that form single-stranded DNA, cause replication forks to stall, and form R-loops. All these features have been observed at the *BRCA1* locus and could be interpreted as consistent with the stress-test hypothesis as it pertains to human *BRCA1*. The hypothesis however predicts these to be features of all *BRCA1* genes for which BRCA1 performs similar functions. Although the human *BRCA1* gene is probably subject to slipped-strand mispairing because of abundant repetitive sequences within its introns, this is not a general feature of mammalian *BRCA1* genes. Whereas *BRCA1* genes of simian primates contain about 50% repetitive sequences, the repetitive fraction is 4% in murine *BRCA1*, 19% in bovine *BRCA1*, and 12% in canine *BRCA1*.

Because BRCA1 participates in pre-mRNA splicing, the stress-test hypothesis also predicts that *BRCA1* will be associated with particular difficulties in its own splicing. Indeed, transcription of *BRCA1* is accompanied by rampant alternative splicing. More than a hundred distinct splicing events have been reported at the *BRCA1* locus and these events are combined into thousands of alternative transcripts, many of them highly unstable (Li et al. 2019). The assembly of functional *BRCA1* mRNA appears to be unusually time-consuming. Knockdown of the U2 snRNP complex reduces transcription of *BRCA1* (Tanikawa et al. 2016) and transient inhibition of the U2 snRNP in triple-negative breast cancers cells resulted in persistent DNA damage and prolonged inhibition of translation of BRCA1 protein (Caggiano et al. 2024). Moreover, BRCA1 interacts with splicing factors in the repair of DNA double-stranded breaks (Lappin et al. 2022; Ozaki et al. 2024). Finally, *RNU2* encodes the U2 snRNA which has a key role in determining the location of 3’ splice sites (Tholen 2024).

A final potential source of epistasis within the *RND2–RNU2* supergene concerns autophagy. NBR1 is a major adapter for targeting cargos to autophagosomes (Odagiri et al. 2021; Rasmussen et al. 2022) and the *NBR2* long noncoding RNA inhibits autophagy (Yang et al. 2024). In most vertebrates, *BRCA1* and *NBR1* share a bidirectional promoter suggesting coordinated regulation of their gene products. Beclin1 is a major mediator of autophagy and *BECN1* (the gene that encodes Beclin1) is located on human chromosome 17 just outside the region of suppressed recombination associated with *BRCA1*. Simultaneous inactivation of *BECN1* and *BRCA1* synergistically increases the sensitivity of ovarian cancers to platinum therapy (Salwa et al. 2021).

*CCDC200* encodes a coiled-coil domain protein that is conserved between sharks and humans but almost nothing is known about its function. CCDC200 proteins of eutherian mammals are glutamine-rich: 37 out of 168 (22%) amino acid residues of human CCDC200 are glutamine, including a stretch of six glutamines in eight residues, and 34 out of 168 (20%) residues of murine CCDC200 are glutamine, including a stretch of five glutamines in seven residues. Polyglutamine tracts in other proteins interact with beclin1 to promote autophagy (Ashkenazi et al. 2017). Beclin1 forms homodimers with itself and heterodimers with other proteins via interactions between coiled-coil domains. The possibility of interactions between Beclin1 and CCDC200 in the regulation of autophagy has not been investigated.

Major unanswered questions are when recombination was first suppressed between *RND2* and *RNU2* and whether the supergene contains strata with different ages of suppressed recombination. Suppressed recombination is believed to promote genetic deterioration because of the reduced effectiveness of purifying selection in eliminating slightly deleterious mutations in the absence of crossing-over (Gutiérrez-Valencia et al. 2021). One signature of this process is an accumulation of transposable elements (Sniegowski and Charlesworth 1994; Stolle et al. 2018). The human *BRCA1* gene contains more than 40% *Alu*-derived sequences, about four times the genomic average, but a high abundance of *Alu* elements is also present in the *BRCA1* genes of macaques (44%) and marmosets (36%). Either recombination has been suppressed since before the divergence of macaques and marmosets or the high abundance of *Alu* elements is a red herring in attempts to date the suppression of recombination. The *BRCA1* loci of GRCh38 (a PEKS haplotype) and CHM13 (an LGRG haplotype) are 99% identical over a distance of 99 kilobases (including near identity for most *Alu* elements). Therefore, the divergence of these haplotypes is much more recent than the high abundance of *Alu* elements at the *BRCA1* locus.

## Conclusions

In modern human populations, near-maximal linkage disequilibrium extends from Jana to Juturna, encompassing *RNU2* and the entire *BRCA1* locus. Among people without recent African descent, there are two major ‘alleles’ of this supergene, which we have labelled PEKS and LGRG, with only rare recombinants between them. European and Asian PEKS haplotypes show evidence of recent expansion to high frequency from a small number of founder haplotypes. There is greater haplotypic diversity among people with recent African ancestry with LEKS the majority haplotype in Africa. In African populations, most haplotypes can be assigned to one of two major haplogroups (PEKS+LEKS and LERG+LGRG). Two patterns require explanation. The first is the maintenance of two major haplogroups in all populations, apparently by some form of balancing selection. The second is the sweep of PEKS to high frequency in non-African populations, displacing LEKS on one side of this polymorphism. A practical consequence of long-range linkage disequilibrium is that, if a trait is correlated with a SNP somewhere within the *RND2–RNU2* supergene, the ‘causal variant’ could reside anywhere within the supergene or the trait could be caused by some combination of variants within the supergene.

The tandem array of *RNU2* repeats is highly homogeneous. Each repeat is associated with a retroviral LTR and the array is immediately adjacent to other retroviral sequences, including the two retroviruses we call ERVK13p and ERVK13d. The *RNU2* array and adjacent retroviral-rich region are notable for their predominance of direct repeats and paucity of inverted repeats that probably predisposes these regions to slipped-strand mispairing. The mechanism of concerted evolution within the *RNU2* array is yet to be determined but may involve unequal exchanges within arrays or involve the retroviral sequences in ‘copy and paste’.

The distal and proximal boundaries of the region of suppressed recombination lie close to the sequences that we call Jana and Juturna. These sequences, together with other repetitive sequences within the supergene, are proposed to form secondary structures that are sources of replicative stress and create barriers to meiotic synapsis. An unusual feature of the *RND2–RNU2* supergene is that recombination appears to be suppressed in supergenic homozygotes as well as heterozygotes whereas inversion polymorphisms typically suppress recombination only in heterozygotes. This unusual region of the human genome deserves focused study.

### Box 1: Evolutionary history of Jana

The short version of Jana can be represented as d–*ACB*–p and the long version as d–*ACBAB*–p (d = distal, p = proximal). The reference sequences of bonobos and chimpanzees possess a d–*ACB*–p arrangement similar to the human short variant. As a benchmark for relative dating, we used evolutionary divergence of a 1.6-kb ERV1 sequence that lies proximal to the Jana structural variant and is present as a single copy in the genomes of humans and great apes. This sequence showed 99% identity between humans and bonobos, 98% identity between humans and gorillas, and 96% identity between humans and orangutans. The two copies of *A* of the human long variant are 99% identical and the two copies of *B* are 93% identical. Based on these comparisons, the duplication of *A* occurred after the human lineage diverged from the lineage of chimpanzees but the duplication of *B* occurred before the last common ancestor of humans and great apes. As further evidence for late duplication of *A*, the two copies of *A* of the human long variant are more similar to each other (99% identity) than either is to the single copy of *A* in bonobos (97–98% identity). This suggests the existence of a third historical arrangements d–*ACBB*–p that was present in an ape–human ancestor and can be referred to as ‘middling Jana’.

The origin of the human long and short variants of Jana appears to have involved slipped-strand mispairing and recombination between two shorter repeated motifs that we will call stutters and stammers. A stutter is a 260-nucleotide sequence, with internal direct repeats, that appears twice in the human short variant and thrice in the human long variant. A stammer is a pair of *Alu* elements that is present five times in the human short variant and seven times in the human short variant. *A* sequences possess two distal stammers and a proximal stutter. *B* sequences possess a distal stutter and a proximal stammer. *C* sequences possess two stutters. In long Jana haplotypes, one stutter performs double duty as part of the proximal *A* sequence and proximal *B* sequence and one stammer performs double duty as part of the proximal *A* sequence and distal *B* sequence. This suggests a scenario in which long Jana arose by slipped-strand mispairing and a double cross-over between an *A* sequence and the duplicated *B* sequences of intermediate Jana, with one of the crossovers occurring between a stammer of *A* and the stammer of the distal *B* sequence and the other occurring between the stutter of *A* and the stutter of the proximal *B* sequence.

*B* sequences can be divided into proximal and distal portions separated by a short internal region containing multiple copies (tics) of a GCCTCTAACC motif. When *B* sequences are partitioned in this way, the proximal portion of the single copy of *B* in the human short variant is 97% identical to the proximal portion of the proximal copy of *B* of human and bonobo long variants but only 91% identical to the distal portion of the proximal copy of *B*. In contrast, the distal portion of the single copy of *B* in the human short variant is 92% identical to the distal portion of the proximal copy of *B* but 97% identical to the distal portion of the distal copy of *B*. Short Jana thus appears to have been derived by recombination between the cluster of tics of a distal copy of *B* and the cluster of tics of a proximal copy of *B* with deletion of intervening sequences. An analysis of the ‘cross-over’ point suggests there may have been independent derivations of short Jana in the human and bonobo lineages but the 97% identity of the *B* sequence of human short Jana to the more similar parts of the proximal and distal B sequences of human long Jana suggests a recombination event before the divergence of African great apes.

### Box 2: Evolutionary history of Janus

The two faces of Janus, together with intervening sequences, have been conserved between humans and baboons, with the distal face of Janus partially overlapping the first coding exon of *CCDC200*. Copies of a sequence homologous to the distal face of Janus and the first exon of *CCDC200* can be found at two locations on marmoset chromosome 5 that are separated by 50 Mb. At both of these locations, ‘half-Janus’ possesses a distal face without a proximal face and is associated with two coding sequences of U2 snRNA. One copy of half-Janus is located near the *ARL4D* gene and the other is associated with the *CCDC200* gene adjacent to *TMEM106A*. The anciently conserved synteny between *VAT1* and *ARL4D* has been disrupted in the marmoset genome with the break occurring near the *CCDC200* first exon and duplicated sequences on both sides of the break. At neither location of half-Janus in the marmoset genome are there sequences related to the retroviral LTRs of the *RNU2* repeats of Old World monkeys, apes and humans.

Between the two faces of Janus, baboons and humans possess sequences that are also present in the *RNU2* repeats of these species. Similar sequences are found in the marmoset genome near *ARL4D* and the U2 snRNA coding sequences. The most parsimonious interpretation of these observations is that the last common ancestor of Old World (catarrhine) and New World (platyrrhine) primates possessed a copy of half-Janus and one or more U2 snRNA coding sequences located between *CCDC200* and *ARL4D*. These sequences were not associated with ERVK13 retroviruses. In the marmoset lineage, a rearrangement separated *ARL4D* and the U2 snRNA coding sequences from *CCDC200, TMEM106A* and *BRCA1*. Before the divergence of Old World monkeys from apes, two events occurred in the human lineage: (a) an inverted duplication created the two faces of Janus from half-Janus and (b) the U2 snRNA coding sequences became associated with ERVK13 retroviruses.

### Box 3: *RNU2*-associated retroviruses

The presence of retroviral sequences adjacent to the human *RNU2* locus was noted by Jones et al. (1994), Liao et al. (1998), and Pavelitz et al. (1999). Orthologs of ERVK13o, ERVK13d, and ERVK13p are present in the reference sequence of bonobos and and orthologs of ERVK13o and ERVK13d, but not ERVK13p, are present in the chimpanzee reference sequence (but the sequence is incomplete in this region). All three retroviruses are absent from the reference sequences of gorillas and orangutans. Although these absences could be interpreted as evidence for recent insertions of ERVK13o and ERVK13p in an ancestor of humans and chimpanzees, the 80% identity of 5’ and 3’ flanking LTRs of both ERVK13o and ERVK13p suggests a substantially older origin because much greater similarity would be expected for a recent insertion.

ERVK13o appears to have descended from the retroviral portion of the *RNU2* repeat of the common ancestor of apes and Old World monkeys because it shows greater than 80% identity to a 5.6-kb sequence of the 11 kb *Papio anubis* (baboon) *RNU2* repeat unit (AF147270). This is consistent with human ERVK13o’s flanking LTRs showing 80% identity to each other and to the LTRs associated with human *RNU2* repeats. The human ERVK13o sequence contains an *Alu* insertion that is not present in the baboon sequence. Its longest open reading frame is *env*-related. Liao et al. (1998) estimated the insertion of ERVK13o as occurring 27 Mya in an immediate common ancestor of Old World monkeys and apes.

Human ERVK13p shows greater than 80% identity to a 10.6-kb sequence adjacent to U2 coding sequences in the reference genome of baboons, including an *Alu* insertion at the same location. Therefore, ERVK13p appears to have already been present near the *RNU2* locus before the divergence of Old World monkeys and hominoids, and has been preserved at this location ever since. ERVK13p possesses substantial open reading frames of reverse transcriptase and integrase domains. The reverse transcriptase and *gag* domains have undergone substantial degeneration. If it were translated, ERVK13p’s integrase could be catalytically active with a conserved integrase domain and a conserved zinc-finger retroviral-genome binding domain, although the integrase has lost its DNA-binding domain for genomic DNA. Notably, the core integrase domain has the conserved DDE amino acid motif, indicative of a catalytically active retroviral integrase, and an intact CCHC structure of a functional zinc finger.

ERVK13d has inserted into the older ERVK13o endogenous retrovirus. The 91% identity between the 5’ and 3’ flanking LTRs of ERVK13d suggest that this was a more recent integration than ERVK13p. Recent integration is also suggested by the absence of *Alu* insertions within ERVK13d. The *gag* domain has not been conserved but the coding sequence of the reverse transcriptase is almost intact (interrupted by a single stop codon) but lacks a functional RNAse H domain. The predicted protein shows close structural similarity to the reverse transcriptase of HIV-1 with an intact N-terminal hydrophobic stretch suitable for nucleotide positioning and 13/13 conserved catalytic core motifs including a YMDD catalytic core (Figure 8). Several sequences in the human genome are highly similar to ERVK13d, including *ERVK13-1* at 16q11.2 and a sequence from the 3’ UTR of *C8orf33* at 8q24.3. *ERVK13-1* was the best match to both ERVK13p and ERVK13d in the gorilla reference sequence. A curious feature of ERVK13d is the insertion of a 1-kb sequence from the 3’-UTR of *DCUN1D3* on human chromosome 16 (conserved at that location from humans to monotremes).

**Figure 8.**
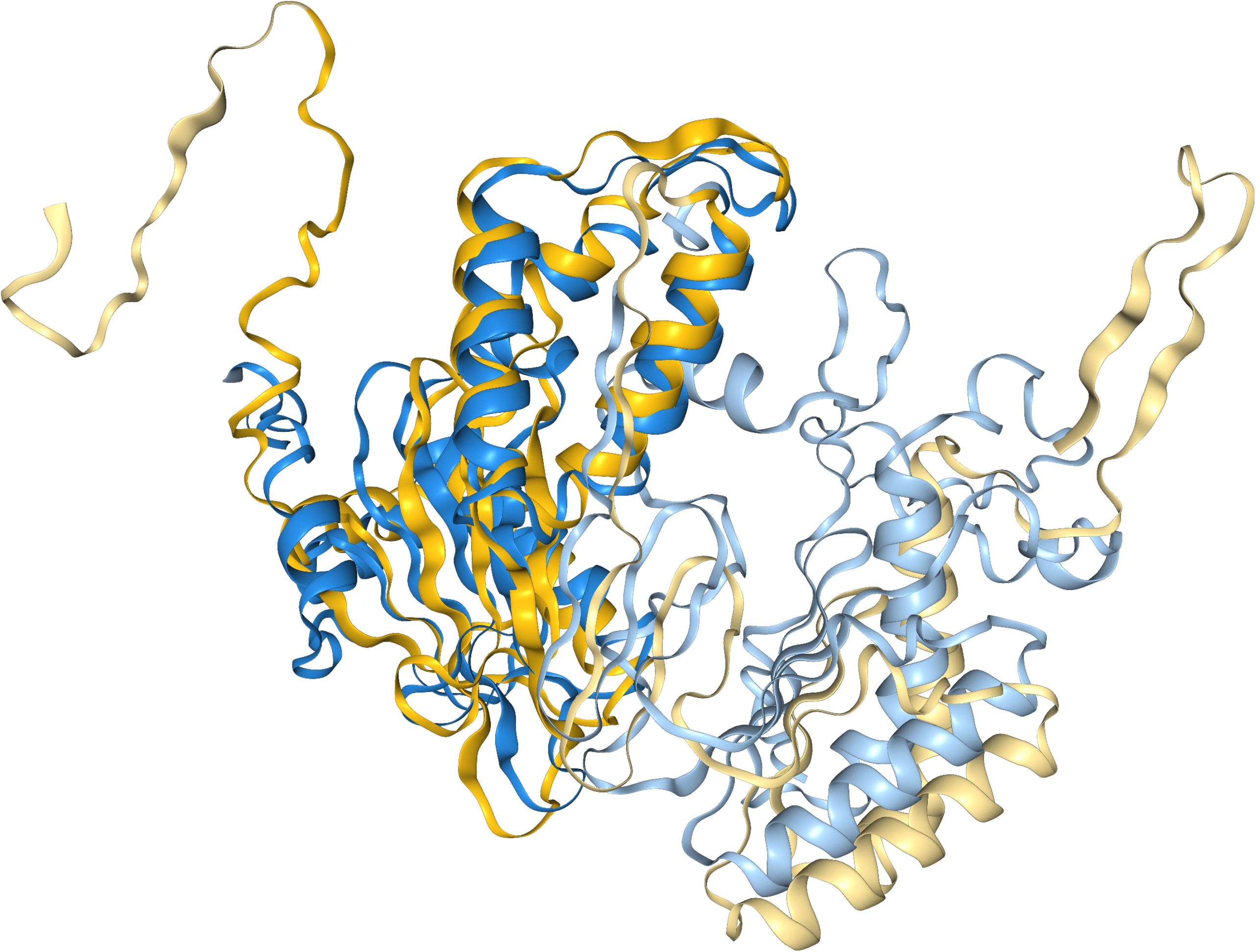
Predicted structure of the putative reverse transciptase of ERVK13d (blue) superimposed on the structure of the HIV-1 reverse transcriptase. Both structures obtained using AlfaFold.

Retroviral proteins can exhibit frameshifts and suppressed stop codons. Moreover, protein domains from one retrovirus can complement domains encoded by another virus. Therefore, experimental evidence is required to determine what functionalities have been retained by ERVK13p and ERVK13d (and whether these functionalities play a role in the concerted evolution of the *RNU2* repeats).

## Methods

All data were obtained from public sources. Phased genetic variants were obtained in vcf format for 2503 individuals with known origin from different populations from the 1000 Genomes Project GRCh38 phased variant dataset (Lowy-Gallego et al., 2019). The telomere-to-telomere (CHM13) was obtained from Nurk et al., 2022 and the GRCh38 assembly from Schneider et al. 2013.

### Identifying the region of Linkage disequilibrium

We identified the region of linkage disequilibrium in two ways. First, we looked at a publicly available genetic map made by the HapMap consortium (International HapMap Consortium, 2007) (made on build 35 (NCBI35; hg17) but lifted to GRCh38), which we obtained from Beagle resources (Browning and Browning, 2007). Secondly, we generated our own linkage disequilibrium maps. Phased genetic variants were obtained in vcf format for 2503 individuals with known ancestry from the 1000 Genomes Project GRCh38 phased variant dataset (Lowy-Gallego et al. 2019), which includes biallelic SNVs and INDELs phased using SHAPEIT2 (Delaneau et al., 2014). Bcftools (Danecek et al. 2021) was used for sample subsetting, genomic region extraction, quality control filtering, monomorphic site removal, and multi-allelic variant normalization. The processed files were converted to binary PLINK format using PLINK2 (Chang et al. 2015) at different levels of minor allele frequency (MAF) filtering (no filter, MAF > 0, 0.01, MAF > 0.05). Pairwise LD statistics (both R^2^ and D’) were calculated using PLINK v1.9 (Purcell et al. 2007) with unlimited window size settings (–ld-window 99999 –ld-window-r2 0). Visualisation was done in Rstudio (Posit team, 2025) using the ‘LDheatmap’ package (Shin et al. 2006).

### Identifying haplotypes

Within this general region, we looked for SNPs that showed strong LD and were present at a minor allele frequence (MAF) greater than 0.25 in Eurasian populations. The non recombining region was then conservatively defined as the region between the most distal and proximal SNPs (rs8071278 and rs12941945; chr17:43041893-43370860 in HG38). Ancestral states were determined based on Chimpanzee (PanTro5 Assembly), Bonobo (Panpan3 assembly) and ancestral reconstruction data (Prado-Martinez et al. 2013). Consensus haplotypes were constructed by selecting the most frequent allele at each position within each haplotype group. To visualize haplotype similarity, we conducted a Principal Component Analysis (PCA). Custom Python v3.8 (Python Software Foundation, 2019) scripts were used. Genotypes were converted to haplotypes using scikit-allel v1.3.0 (Miles and Harding 2017). PCA analyses were implemented using scikit-learn (Pedregosa et al. 2011) and NumPy (Harris et al. 2020), with visualization performed through Matplotlib (Hunter 2007).

### Haplotype ancestries

We obtained variant data for ancient 36 European humans (30,000–40,000 years old) and seven Neanderthals through personal contact with David Reich’s lab. Variant calling was performed using the bcftools “mpileup” and “call” with multiallelic calling mode (Li, 2011; Danecek et al. 2021). The DNA reads were trimmed on both ends to remove damaged artifacts, once with 5 bp (conservative) and once with 2 bp (less conservative, but less information lost). Quality filtering was applied to remove low-confidence calls, specifically sites with read depth <4 or variant quality score <20, which were marked as missing data. Coordinate conversion between GRCh37 and GRCh38 was performed using CrossMap (Zhao et al. 2014). Using the 2bp-trimmed variant calls and considering the uncertainty on the read ends and also considering potential post-mortem damage (C to T mutations on 5’-end and G to A on the reverse complement 3’-end), we manually assigned haplocodes based on the four tagging SNPs to the ancient human and Neanderthal sequences. Pairwise genetic distances (Nei 1972) were calculated between the previously made consensus haplotypes for PEKS, LEKS, LGRG and LERG, a consensus *Pan* haplotype (most frequent alleles for twenty-five chimpanzees and thirteen bonobos), a randomly chosen Neanderthal and a randomly chosen homozygous PEKS-like ancient human sample.

### Structural variation

GRCh38 (Schneider et al. 2017) reference has a PEKS haplotype and CHM13 (Nurk et al., 2022) assembly has an LGRG haplotype. We performed BLASTN alignment (Altschul et al. 1990) of the *RND2–RNU2* region between the GRCh38 and CHM13 reference genome assemblies, as these represent different haplotypes (PEKS and LGRG respectively).

We also obtained structural variant data in vcf format from the 1000 Genomes ONT Vienna collection (Schloissnig et al. 2025). We isolated fifteen structural variants within our region of interest and then looked for variants linked to PEKS, LGRG and LEKS haplotypes.

### Phenotypic effects and expression

Investigating the phenotypic effects of the different haplotypes was done in four steps. First, a GWAS database search was performed on the SNPs that differ between the LGRG and PEKS consensus haplotypes. We looked up these SNPs in the GWAS catalog (MacArthur et al. 2017) and selected those with pvalue < 5·10^8^. This provided fifteen SNPs associated with GWAS hits (Table S3). Next, We identified differences in gene expression between the LGRG and PEKS haplotypes based on and GTEx RNA-Seq data (GTEx Consortium et al. 2020). We identified all expression quantitative trait loci (eQTL) hits that were significantly correlated with one of the previously identified PEKS/LGRG haplogroup specific SNPs. We performed a similar analysis on splice quantitative trait loci (sQTL) data from the same source. Haplogroup specific SNPs were also looked up in clinical data obtained from the Open Targets Platform (Carvalho-Silva et al. 2019). Finally, variant consequences for the haplogroup specific SNPs were using the Ensembl Variant Effect Predictor (McLaren et al. 2016). Polyphen (Adzhubei et al. 2010) was used to predict whether a SNP was benign or harmful.

### Site frequency spectrum and neutrality statistics

Custom python scripts were used for a comprehensive site frequency spectrum (SFS) analysis in order to characterise allele frequency distributions and provide neutrality statistics across different (super)populations and haplogroups. VCF files were processed to isolate individual haplotypes of the *RND2*–*RNU2* region from specific haplogroups and/or (super)populations. This was done for all haplotypes combined and also for isolated haplogroups (PEKS, LGRG and LEKS only). Statistics were calculated if at least ten haplotypes are present in the subgroup. Ancestral states were inferred using the *Pan troglodytes ellioti* and *Pan paniscus* genomes. Unfolded site frequency spectra were made using these ancestral states and Tajima’s D and Fay & Wu’s H were calculated (Tajima 1989; Fu and Li 1993; Fu 1997; Fay and Wu 2000).

### Genomic architecture and replicative stress

The genomic sequence of the *RND2–RNU2* region of human chromosome 17 was investigated in detail using the telomere-to-telomere assembly (CHM13) because of problems with GRCh38 in this region. We identified several regions of interest using BLASTN alignments (Altschul et al. 1990) of CHM13 to itself, to GRCh38, and to the bonobo reference. Comparisons with other primates were performed using the UCSC genome browser (Kent et al. 2002). Similarity scores between sequences were taken from BLAST alignment percentage identity scores (Altschul et al. 1990). We used Unafold (Markham and Zuker 2008) to look at predicted structure formation and RepeatMasker (Smit et al., 2013) to identify repeats.

### Retroviral open reading frame prediction

All open reading frames were manually annotated between chr17:43,307,126 and chr17:43,383,785 of the GRCh38 assembly. Functional domains of open reading frames longer than 100 amino acids were annotated using NCBI conserved domain search tool (CD-Search). Retroviral core catalytic residues were aligned and identified using this tool. Open reading frames with predicted functional domains were folded using AlphaFold 3. All predicted structures were aligned to curated PDBs of retroviral proteins using FoldSeeker. Intact retroviral-like proteins were aligned using ChimeraX protein viewer to ensure catalytic residue positioning.

## Acknowledgments

We thank Sergei Mirkin for discussion of H-DNA structures and thank Shop Mallick and David Reich for help with access to archaic and ancient human DNA sequences.

